# Categorical Bayes filtering for computational phenotyping in adaptive learning

**DOI:** 10.64898/2026.05.14.725268

**Authors:** Junxi Chen, Payam Piray

**Author notes:** Now at New York University (Junxi Chen).

## Abstract

Adaptive learning depends on inferring the sources of uncertainty in the environment. Simultaneous estimation of two such sources, volatility and stochasticity, is difficult, but critical for adaptive learning. The two require opposite adjustments of the learning rate, yet both inflate the variance of experienced outcomes, and no closed-form solution exists for dissociating them. Particle filters are the standard method for this joint inference, but their likelihood is stochastic and their objective is not differentiable, so fitting requires derivative-free methods that add estimator noise. This is a serious problem for individual-difference analyses, because estimator noise attenuates any correlation between fitted parameters and other measures. Here we introduce the Categorical Bayes Filter (CBF), a deterministic alternative. The CBF keeps the conditional structure of recent particle-filter models, but replaces the stochastic outer layer with a categorical distribution over a quantile grid, parameterized by differentiable Beta quantile functions. Fitting is therefore evidence maximization with an exact, deterministic, and differentiable marginal likelihood. In a large-scale study of a volatility-stochasticity task, the CBF outperformed a state-of-the-art particle filter in predicting trial-by-trial empirical learning rates, its dispersion parameters captured volatility-blind and stochasticity-blind participants, and its trial-by-trial ambiguity predicted response times. Fitted CBF parameters also predicted psychiatric symptoms in the direction previously hypothesized. More broadly, the CBF applies to any particle-filter model with continuous latent parameters, and makes individual-level analyses with such models practical in cognitive modeling and computational psychiatry.

## 1 Introduction

Adaptive learning requires distinguishing two sources of uncertainty: *volatility*, changes in the latent state of the environment, and *stochasticity*, moment-to-moment noise in observations. The two demand opposite responses: a volatile environment calls for a higher learning rate, while a stochastic one calls for a lower one. Disentangling them is computationally hard because both inflate the variance of observed outcomes; correctly attributing surprise to one versus the other is an explaining-away problem in which the two estimates are mutually coupled [1]. This structure rules out the standard hierarchical Bayesian approximations that fix one source while inferring the other [2–4]: they cannot represent the competition that defines the inference. Capturing it requires joint inference over both noise sources, and that is where the practical difficulties begin.

Particle filters are the natural tool for this kind of joint inference, and they have been deployed across a range of cognitive models, including category learning [5], conditioning [6], causal inference [7], change detection [8], sentence processing [9], attention [10], and joint volatility-stochasticity inference [11, 12]. Among the latter, the Rao-Blackwellized particle filter (RBPF) is used to confine the particles to the low-dimensional noise-parameter space by integrating out the latent state via a conditional Kalman filter, recovering known signatures of human adaptive learning [11, 12]. Yet across all of these applications the same practical problems recur, and they all stem from the stochastic outer layer. The likelihood is both stochastic, evaluating the same model on the same data twice yields different numbers, and non-differentiable with respect to the generative parameters. The field has compensated with derivative-free optimizers, but these inherit the stochasticity of the likelihood, provide no second-order information, and lack the gradient access that supports modern Bayesian and empirical-Bayes fitting workflows. The downstream consequences are familiar: model comparison is fragile because likelihoods drift across runs, parameters are not reliably recoverable from finite data, and trial-by-trial belief trajectories are obscured by sampling variance. The cost is highest for cognitive modeling more broadly, where the standard paradigm fits a separate copy of the model to each subject (human or animal) from a few hundred trials of behavioral data. Trait-relevant and individual-difference effects in this literature are intrinsically modest, and any additional measurement noise from the estimator suppresses them further.

This paper introduces the Categorical Bayes Filter (CBF), a deterministic alternative that preserves the Rao-Blackwellization insight, separating tractable conditional inference from intractable parameter inference, but replaces the stochastic outer layer with a categorical distribution over a quantile grid. The grid is parameterized through differentiable Beta quantile functions, which makes the marginal likelihood both deterministic and a smooth function of its parameters; gradient-based optimization, hierarchical estimation, and Laplace approximation all become available, and refit-to-refit variability is eliminated by construction. The procedure factors into two phases: exact discrete Bayesian inference under the prior, and gradient-based optimization of the prior to maximize the resulting evidence. The representational strategy borrows from distributional reinforcement learning [13]: anchoring the support and updating only probability masses yields stability that drifting particles cannot match.

On a 2 × 2 factorial volatility–stochasticity task with *N* = 643 participants [11], the CBF (i) reveals a cross-over phenotyping pattern in fitted dispersion parameters, with volatility-blind subjects showing narrow volatility grids while stochasticity-blind subjects show narrow stochasticity grids, an effect that the RBPF on the same data does not produce; (ii) recovers its parameters reliably from synthetic data; (iii) outperforms the RBPF on trial-level prediction of human learning rates while doing so deterministically; (iv) produces a trial-by-trial ambiguity signal that predicts human response times; (v) is preferred over the RBPF in formal random-effects Bayesian model selection; and (vi) links the stochasticity dispersion parameter to psychiatric dimensions. Beyond this task, CBF achieves stronger parameter recovery than an independent RBPF implementation in a social latent-cause inference task [14], demonstrating its applicability across distinct latent structures and its potential for reliable individual-level inference across cognitive domains.

Code is available at https://github.com/piraylab/CBF.

## 2 Related work

### Adaptive learning under uncertainty

A long line of work in cognitive neuroscience has formalized adaptive learning as Bayesian inference over latent environmental structure, with the central question being how to update beliefs when the reliability of incoming information is itself uncertain. Early accounts emphasized a single source of uncertainty governing the learning rate, typically environmental volatility [2, 3] or expected and unexpected forms of noise [15, 16]. Closed-form approximations such as the Hierarchical Gaussian Filter [3] and delta-rule schemes with adaptive learning rates [17] made these models tractable enough for fitting to behavioral data, and they have been used extensively to study individual differences in learning, including in clinical populations [18, 19]. More recent theoretical work has argued that volatility and stochasticity compete to explain experienced variance, and that this competition is the substantive content of the inference problem [1, 11]. Approximations that fix one noise source while inferring the other cannot represent this competition, and so cannot reproduce the behavioral signatures it predicts. The present work takes the dual-noise framework as given and focuses on the inference machinery required to fit it at scale.

### Particle filters in cognitive modeling

Across cognitive applications of particle filters, the same set of practical limitations recurs. The marginal likelihood is stochastic, in the sense that two evaluations of the same model on the same data return different numbers, and it is non-differentiable with respect to the generative parameters. Fitting therefore relies on derivative-free optimizers run on noisy objectives, which means parameter estimates inherit variance from both the likelihood and the optimizer. The Rao-Blackwellized particle filter [20] is particularly natural for our application of joint volatility-stochasticity inference [1, 11], since the latent state can be integrated out analytically conditional on the noise parameters; but the resulting marginal likelihood over the noise parameters remains stochastic and non-differentiable, inheriting the same limitations. Particle filter-based cognitive models have largely served as theoretical demonstrations rather than practical inferential tools, and quantitative individual-difference analyses with them remain rare.

### Differentiable and variational extensions of sequential Monte Carlo

A separate line of work in machine learning has sought to make particle filters compatible with gradient-based optimization. Variational sequential Monte Carlo methods [21–23] treat the SMC marginal likelihood estimator as a stochastic lower bound and optimize it using reparameterization or score-function gradients. Differentiable particle filters [24, 25] replace discrete resampling with differentiable surrogates such as soft resampling or optimal-transport-based couplings. The marginal likelihood in these methods remains stochastic, so refit-to-refit variability is reduced but not eliminated. [26] (DPVI) is the closest prior work in spirit, replacing stochastic resampling with deterministic search over particle locations and weights. DPVI is built for high-dimensional combinatorial latent structure such as Dirichlet process mixtures and infinite HMMs, where the discreteness of the structure precludes gradient-based optimization. The CBF addresses a different regime within the same family: low-dimensional bounded continuous parameter spaces, where the support can be parameterized analytically through differentiable quantile functions and the optimization becomes a smooth gradient ascent over a small number of shape parameters. The two methods share the deterministic-likelihood goal but make opposite choices about how the particle support is optimized: search for DPVI, gradient ascent for CBF.

### Distributional reinforcement learning and grid-based filtering

The fixed-support categorical representation in the CBF was inspired by distributional reinforcement learning. The C51 algorithm [13] demonstrated that maintaining a categorical distribution over a fixed set of return atoms achieves substantial improvements over conventional value-based reinforcement learning methods, and subsequent work [27, 28] generalized this insight to quantile and implicit representations. The CBF transposes this representational choice to Bayesian inference over parameters: rather than letting particles drift through the parameter space, atoms are placed deterministically and only their probability masses are updated. The connection is one of design pattern rather than mathematical equivalence; what carries over is the intuition that fixing the support removes a source of optimization instability. Independently, grid-based and point-mass filters have a long history in nonlinear filtering and tracking [29, 30], where they served as practical alternatives to extended Kalman filters before the rise of particle methods. The CBF is a modern instance of this older idea, augmented in two specific ways: the grid is parameterized by differentiable quantile functions of a flexible bounded family, and the placement of the grid is itself optimized by gradient ascent on the exact marginal likelihood.

## 3 Method

This section develops the Categorical Bayes Filter (CBF) as a deterministic and fully differentiable alternative to particle-filter inference. The framework is presented for a generic sequential inference problem and then instantiated for the joint inference of volatility and stochasticity.

### 3.1 General setting

Consider a sequential inference problem in which an agent observes outcomes *o*_1_, … , *o*_*T*_ generated by a latent process governed by a static parameter vector *θ* ∈ Θ ⊆ ℝ^*D*^. The agent does not know *θ* and must infer it online. Conditioned on *θ*, a procedure is assumed to produce the predictive likelihood of each new observation,

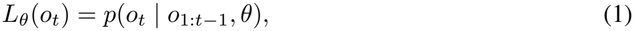

where the procedure may be an analytic filter (such as a Kalman filter that maintains sufficient statistics 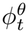 and integrates out latent states in closed form), a numerical integration, or a simulation-based approximation. The CBF framework is agnostic to its internals. The true marginal likelihood,

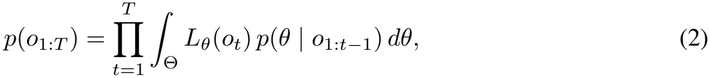

is generally intractable, and standard particle-filter approaches approximate it by stochastic sampling over *θ*.

### 3.2 Categorical Bayes filter

The CBF resolves the optimization bottlenecks of the Rao-Blackwellized Particle Filter (RBPF) by replacing stochastic particles with a fixed grid of *K* atoms *G* = {*θ*_1_, … , *θ*_*K*_} ⊂ Θ, maintaining a categorical distribution over them. Instead of approximating a continuous prior via Monte Carlo sampling, we treat the generative model as possessing a parameterized discrete prior over *G*. This eliminates the non-differentiable resampling step of standard particle filters, reducing inference to a fully differentiable two-phase procedure.

#### Phase 1: Exact discrete inference

Initialize the prior weights uniformly, *w*_*k*,0_ = 1*/K*. On each trial *t*, the weights are updated via Bayes’ rule on the discrete support,

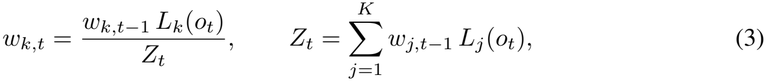

while per-atom internal states advance via 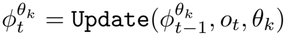.

Because the support is finite, the procedure constitutes exact Bayesian inference under the assumed discrete prior. The sequence of log-normalizing constants log *Z*_*t*_ serves a dual purpose: they ensure the posterior normalizes to 1 at each step, and their sum yields the exact log marginal likelihood of the data under the discrete grid *G*:

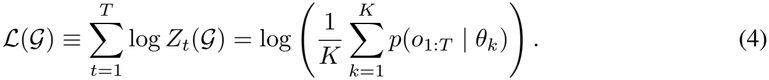

This identity follows by induction. At *t* = 1, the normalizer is the marginal probability of the first observation: 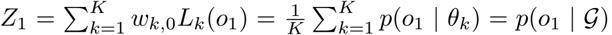. At *t* − 1, assuming the posterior weights are exact, *w*_*k,t* −1_ = *p*(*θ*_*k*_ | *o*_1:*t* − 1_, *G*), the normalizer at *t* expands via the product rule:

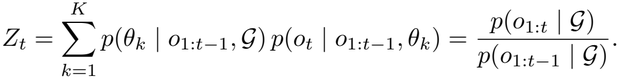

Summing the log-normalizers,

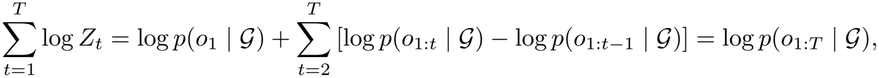

where intermediate terms cancel.

#### Phase 2: Differentiable grid optimization

To ensure the discrete approximation accurately captures the high-density regions of the true continuous parameter space, the grid is parameterized by *ψ* via the quantile function of a flexible bounded distribution. For each dimension *i, K*_*i*_ atom locations are placed at equally spaced quantiles 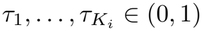:

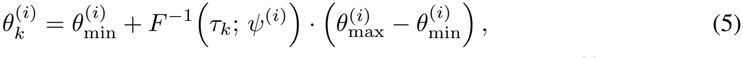

where *F* ^−1^(·; *ψ*^(*i*)^) is the quantile function of a Beta distribution with parameters *ψ*^(*i*)^. The full grid *G*(*ψ*) is the Cartesian product across dimensions.

The optimization phase maximizes the exact discrete marginal likelihood via evidence maximization:

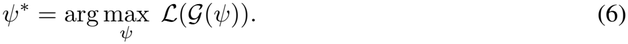

Because the Beta quantile function is differentiable in *ψ* and *Z*_*t*_ is smooth with respect to the atom locations, ℒ (*G* (*ψ*)) is fully differentiable end-to-end, permitting direct gradient ascent on the exact log-evidence.

### 3.3 Application to joint inference of volatility and stochasticity

The framework is instantiated for joint inference of volatility *v* and stochasticity *s* in a Gaussian state-space model.

#### Parameters and conditional filter

The parameter vector is *θ* = (*v, s*) ∈ [*v*_min_, *v*_max_] × [*s*_min_, *s*_max_], so *D* = 2. For each atom *θ*_*k*_ = (*v*_*k*_, *s*_*l*_), the conditional filter is a Kalman filter with process noise *v*_*k*_ and observation noise *s*_*l*_. The filter state 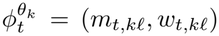 holds the posterior mean and variance of the latent state *x*_*t*_, and the predictive likelihood is

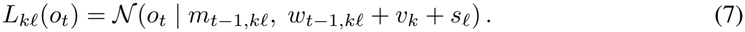

#### Beta parameterization

The Beta distribution is parameterized by its mean *µ* ∈ (0, 1) and dispersion *η* ∈ (0, 0.5), with shape parameters *α* = *µ/η* and *β* = (1 − *µ*)*/η*. The dispersion is the reciprocal of the concentration *α* + *β* = 1*/η* and is monotonic in the variance: small *η* corresponds to a tightly concentrated Beta, large *η* to a flat one. The bound *η <* 0.5 ensures *α* + *β >* 2, keeping the Beta away from U-shaped configurations and the quantile function numerically well-behaved on the interior quantiles used to place atoms.

#### Grid construction

With *M* atoms per dimension (*K* = *M* ^2^), the grid parameters are *ψ* = (*µ*_*v*_, *η*_*v*_, *µ*_*s*_, *η*_*s*_). Atom locations specialize Eq. 5 to the Beta family:

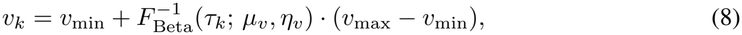

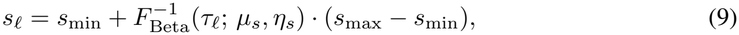

with *τ*_1_, … , *τ*_*M*_ linearly spaced in [0.005, 0.995].

## 4 Results

We evaluated the CBF on a previously published 2 × 2 factorial volatility-stochasticity task with *N* = 643 human participants [11]. The task crosses two levels of environmental volatility (stable, volatile) with two levels of observation stochasticity (low, high), yielding four blocks of 50 trials per subject. The same dataset was used to fit the RBPF, providing a direct head-to-head comparison. To illustrate the model’s behavior, Fig. 1 shows tracking of distinct volatility and stochasticity levels in a synthetic 2 × 2 design.

**Figure 1:**
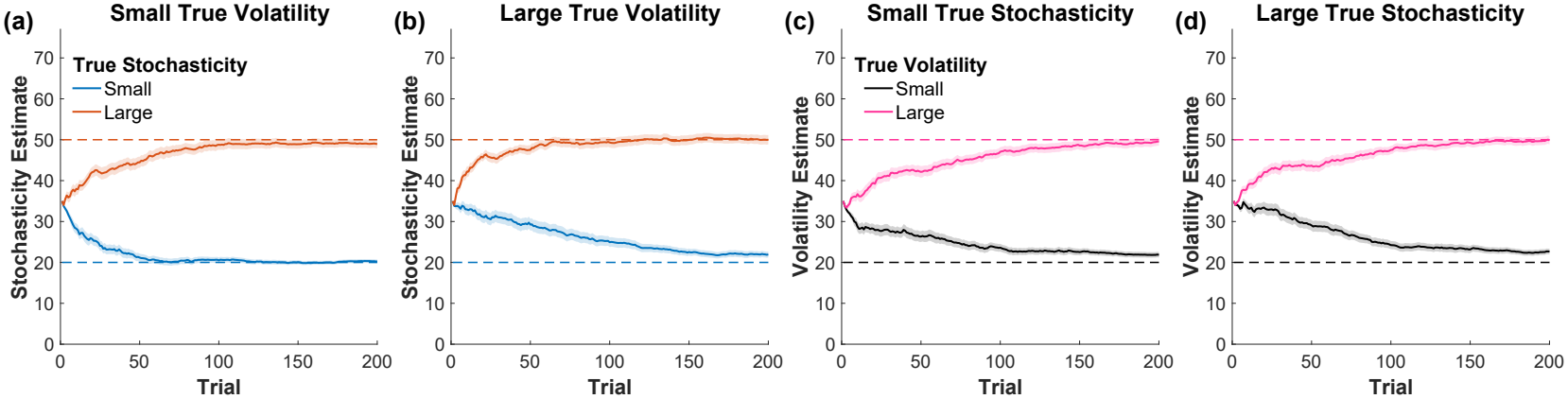
CBF dynamics on a synthetic. 2 × 2 **design. (a-d)** Tracking of true volatility and stochasticity across four blocks of a synthetic 2 × 2 factorial design (200 trials per block; true values of volatility and stochasticity chosen for illustration). Each panel shows the model’s mean posterior estimate of one noise dimension when the other is held at one of two levels: **(a, b)** stochasticity estimates under small (a) and large (b) true volatility; **(c, d)** volatility estimates under small (c) and large (d) true stochasticity. Lines and shaded regions show mean and standard error of the mean across 50 simulations. Dashed lines indicate true values.

### 4.1 Parameter recovery

We first evaluated whether the CBF can recover its parameters from synthetic data. Synthetic datasets were generated from the CBF generative model with parameters drawn uniformly across the admissible range, matching the task structure of the empirical experiment (4 blocks of 50 trials per simulated agent). Models were then fit using the same procedure later applied to human data, including weakly informative Gaussian priors on the unconstrained parameters. Recovery was high across all four parameters (Spearman *r >* 0.98; Fig. 2), with mild shrinkage toward the center of the parameter space for parameters near the boundary of the simulation range. This shrinkage is conservative: it biases recovered parameters toward the center of the parameter space rather than amplifying individual differences.

**Figure 2:**
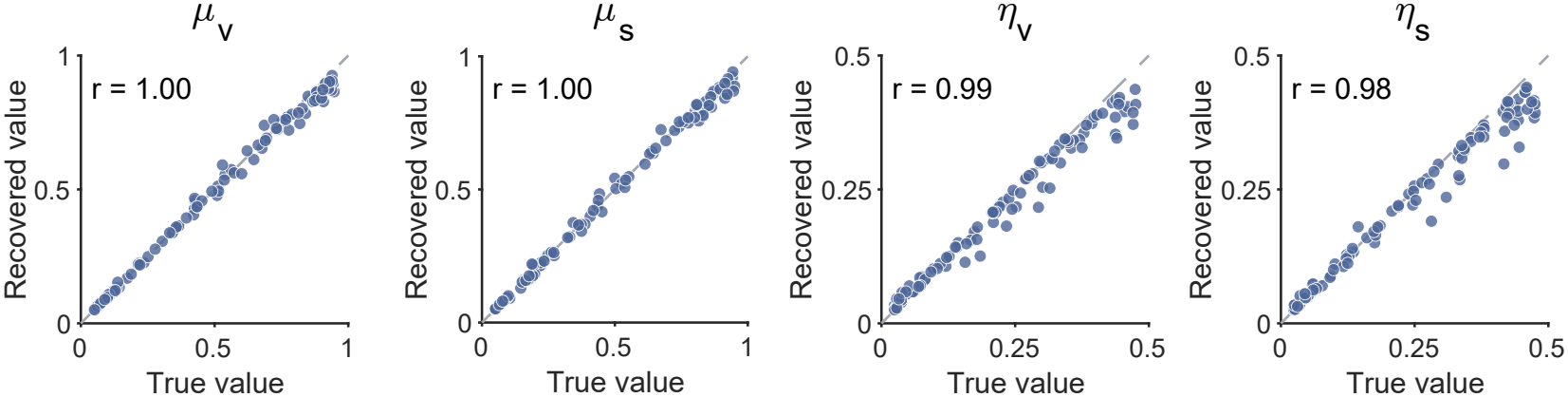
Parameter recovery for the CBF on synthetic data. Scatterplots of true versus recovered values for each of the four CBF parameters: *µ*_*v*_, *µ*_*s*_, *η*_*v*_, and *η*_*s*_. Synthetic datasets were generated from the CBF generative model with parameters drawn uniformly across the admissible range, then fit using the same procedure applied to human data, including the weakly informative Gaussian prior on the unconstrained parameterization. Recovery is high across all four parameters (Spearman *r >* 0.98). Mild shrinkage toward the center of the parameter space is visible for parameters near the boundary of the simulation range; this is a conservative bias that diminishes recovered individual differences rather than amplifying them.

Recovery of the RBPF’s volatility and stochasticity update rates on synthetic data (generated by the RBPF model itself) is substantially noisier: the correlation between true and recovered values is *r* = 0.69 for the volatility update rate and *r* = 0.67 for the stochasticity update rate. The gap reflects the same underlying source of noise that limits RBPF cross-seed reliability on real data: the stochastic likelihood and derivative-free fitting procedure together produce parameter estimates that incompletely identify the underlying subject, even when the data are generated from the model itself.

### 4.2 Computational phenotyping

#### Theoretical background and prediction

Volatility and stochasticity make opposite demands on the learning rate: a volatile environment renders prior beliefs obsolete and calls for rapid updating, while a stochastic environment renders new observations unreliable and calls for caution. Yet both inflate the experienced variance, so the learner has no direct cue distinguishing them; correctly attributing surprise to one source requires explaining away the other [1]. When the prior over one source is too narrow to support this competition, the inference fails systematically: surplus surprise is routed entirely to whichever dimension the prior can represent. A learner with a narrow stochasticity prior therefore sees the world as more volatile than it is, and a learner with a narrow volatility prior sees it as more stochastic. The behavioral consequence is not impaired but reversed learning-rate modulation. Behavioral validation of this framework in a large online sample found that, on average, participants showed the predicted pattern: a positive main effect of true volatility on the learning rate and a negative main effect of true stochasticity [11]. However, there was substantial individual variation around these group-level means, with sizable fractions of participants showing reversed sensitivity to one of the two noise sources while remaining adaptive to the other.

#### Behavioral classification of subjects

The task is designed so that learning rates can be estimated model-agnostically: on each trial, participants report a prediction (by positioning a bucket to catch a bag of coins, the observation), and the trial-by-trial learning rate can be recovered as the regression coefficient of the prediction update on the prediction error [11]. True volatility and true stochasticity are included as interaction terms with the prediction error, yielding two critical coefficients per subject: the main effects of volatility and stochasticity on the learning rate. Adaptive subjects show a positive volatility main effect (learning rate increases with volatility) and a negative stochasticity main effect (learning rate decreases with stochasticity), in line with the normative prediction. Across the *N* = 643 participants, both effects were highly significant in the predicted directions, although individual subjects varied substantially around these averages [11].

To isolate subjects whose behavior is selectively pathological on one noise dimension while remaining clearly adaptive on the other, we used the sample-mean main effects as thresholds. Let 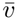 denote the mean volatility main effect across all 643 subjects (a positive value) and 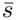 the mean stochasticity main effect (a negative value). Subjects were classified as *volatility-blind* when their volatility main effect was below 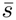 (i.e., strongly reversed) while their stochasticity main effect was also below 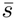 (i.e., adaptive sensitivity on stochasticity stronger than the population average). Symmetrically, subjects were classified as *stochasticity-blind* when their stochasticity main effect was above 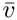 (strongly reversed) while their volatility main effect was also above 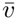 (adaptive sensitivity on volatility stronger than the population average). These criteria yielded 76 volatility-blind and 79 stochasticity-blind subjects. The defining feature of these two groups is that the maladaptation is specific to one dimension while sensitivity on the other dimension is preserved at a level above the population average, making them reliable phenotypes rather than diffuse low-performers. The classification is purely behavioral and made before any model is fit.

#### The cross-over pattern in CBF parameters

The CBF framework provides a direct test of what behavioral blindness to one noise dimension means at the level of the inferred prior. A subject whose behavior fails to track variation in one noise source must, by the logic of the framework, hold a prior that is too narrow to represent that variation. The prediction is therefore straightforward: volatility-blind subjects should have low *η*_*v*_ (a narrow volatility prior), and stochasticity-blind subjects should have low *η*_*s*_ (a narrow stochasticity prior), with each group showing typical dispersion on the dimension they are not blind to. This is exactly the pattern observed in the data (Fig. 3, left): volatility-blind subjects had significantly lower *η*_*v*_ than stochasticity-blind subjects (*t*(153) = − 5.23, *p <* 0.001), while stochasticity-blind subjects had significantly lower *η*_*s*_ (*t*(153) = − 10.01, *p <* 0.001). The two contrasts together produce the cross-over predicted by the framework: each group is concentrated on its own blind dimension and at typical dispersion on the other.

**Figure 3:**
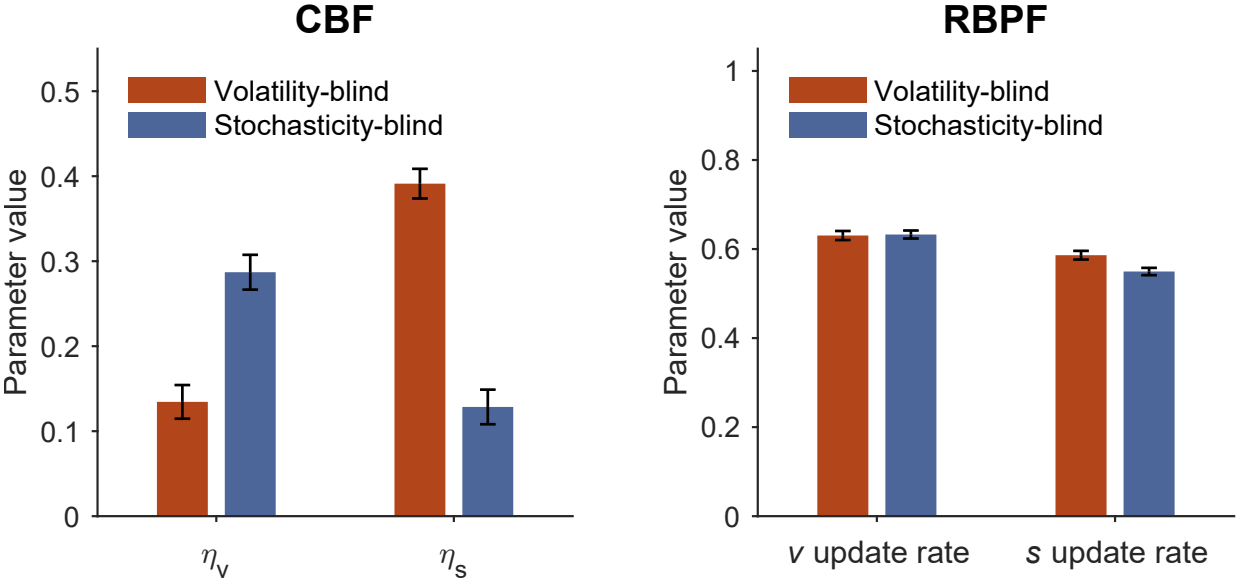
Cross-over phenotyping in human subjects: CBF dispersion parameters reveal dimension-specific pathology, while RBPF parameters do not. Mean fitted CBF dispersion parameters *η*_*v*_ and *η*_*s*_ for volatility-blind subjects (*n* = 76) and stochasticity-blind subjects (*n* = 79); each group shows a narrow prior on its blind dimension (low dispersion) and typical dispersion on the other, producing a cross-over. The same comparison applied to the RBPF volatility and stochasticity update rates (denoted *µ*_*v*_, *µ*_*s*_ in [11]) does not produce the cross-over. Y-axis scales differ between panels because the parameters have different admissible ranges. Error bars denote standard error across subjects.

The cross-over results were robust to grid configuration and held without any grouping: both group contrasts survived re-fitting across five grid-resolution and support settings, and across all *N* = 643 participants, *η*_*v*_ and *η*_*s*_ still predicted their corresponding behavioral main effects after controlling for *µ*_*v*_ and *µ*_*s*_ (Appendices E and G).

#### Comparison to RBPF

The same analysis was applied to the RBPF’s volatility and stochasticity update rate parameters (taken from the public code repository associated with [11]), which capture the agent’s beliefs about how volatility and stochasticity vary over trials. The RBPF parameters do not produce the cross-over (Fig. 3, right). This is not a consequence of casual fitting: the RBPF was fit using a protocol combining gradient-free Bayesian optimization with a regularized objective averaged over ten random seeds per parameter evaluation. The averaging and regularization were designed precisely to mitigate the non-determinism inherent in particle filtering, and the resulting fits are about as stable as the method allows. Even so, residual variability is unavoidable: refitting the RBPF to the same human subjects with two different top-level seeds yields fitted parameters that correlate at *r* = 0.92 and 0.83 across seeds for the volatility and stochasticity update rate parameters respectively. By the logic of measurement attenuation, this level of refit noise reduces any observed correlation between fitted parameters and external measures, which is consequential in a literature where trait-parameter and behavior-parameter associations are typically of small to medium effect size. The CBF avoids this attenuation by construction, since its dispersion parameters are deterministic and remain well-defined across the parameter range; we return to why this matters specifically for the pathological subjects in the discussion.

#### External validation against psychiatric symptoms

We further examined whether the fitted parameters were associated with psychiatric symptom measures collected after the learning task in a subset of participants (*n* = 401). The framework predicts that low *η*_*s*_ should accompany the intolerance of uncertainty and chronic worry characteristic of anxiety [1], and this was the case: *η*_*s*_ was negatively associated with a standardized worry-minus-depression contrast (*r* = − 0.157, *p* = 0.0016), with lower *η*_*s*_ corresponding to greater worry relative to depressed mood. Note the contrast was constructed after examining the relationship between worry and depression scores (Appendix H).

### 4.3 Trial-level prediction of learning rates

A standard test of adaptive-learning models is whether the model’s trial-by-trial learning rate tracks the empirical learning rate computed from each subject’s behavior. The CBF achieved a rank correlation of *r* = 0.866 with the across-subject average empirical learning rates of the *N* = 643 participants, exceeding the RBPF’s *r* = 0.800 on the same data. A more diagnostic test asks whether the model captures how the learning rate is modulated by each of the two noise sources it is built to dissociate. We computed the volatility and stochasticity main effects on the learning rate separately for the model-derived and empirical trajectories, and correlated the per-subject coefficients across all participants. The CBF tracked the volatility main effect with Pearson *r* = 0.52 and the stochasticity main effect at *r* = 0.61, substantially exceeding the RBPF’s *r* = 0.34 and *r* = 0.36. To contextualize these correlations, we estimated the test-retest reliability of the empirical main effects via Spearman-Brown-corrected odd-even split-half correlation: *r*_*xx*_ = 0.70 for the volatility effect and *r*_*xx*_ = 0.78 for the stochasticity effect, both high for individual-difference measures in cognitive tasks. The corresponding ceiling on any model-behavior correlation is 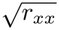 or about 0.84 and 0.88. CBF therefore reaches approximately two-thirds of the achievable correlation on both dimensions (62% on volatility, 69% on stochasticity), compared to roughly 40% for the RBPF. The deterministic representation does not sacrifice fidelity to behavior; if anything, it gains it on the dimensions the model is built to explain. Fitting is also achieved at a fraction of the cost: per-subject fitting takes seconds rather than the hour required for the RBPF protocol, with a deterministic likelihood that supports gradient-based hierarchical estimation at scale.

### 4.4 Trial-by-trial prediction of response times

The CBF maintains a deterministic categorical posterior over the noise-parameter grid on every trial, and the standard deviation of predictive likelihoods across atoms provides a trial-by-trial measure of *computational ambiguity*: the degree to which current evidence fails to discriminate between competing noise hypotheses. When likelihoods are concentrated on a small subset of atoms, ambiguity is low; when likelihoods are spread, ambiguity is high. This quantity is a deterministic function of the data given a fitted model.

Trial-level ambiguity predicted human response times: trials with higher ambiguity (lower likelihood SD, i.e., likelihoods spread across many atoms) were associated with slower responses (Table 2; per-subject linear regression, *β* = − 8.93, *t* = − 4.58, *p <* 0.001, controlling for mean predictive likelihood, estimated volatility and stochasticity, and squared prediction error). Response times were not used in fitting the model. The relationship is therefore a held-out prediction from a model fit to choice data alone, and it provides a process-level signature of the explaining-away computation.

### 4.5 Bayesian model selection

A random-effects Bayesian model selection [31, 32] was performed across the *N* = 643 subjects, treating CBF and RBPF as competing models of trial-by-trial choice behavior. The CBF was favored over the RBPF with a protected exceedance probability of 1.0000 and an expected model frequency of 0.6471 versus 0.3529 for the RBPF (Table 3).

### 4.6 Generalization to a social latent-cause task

To assess generality beyond the 2 × 2 factorial volatility-stochasticity task, we compared parameter recovery in a social latent-cause inference task [14] using an independent RBPF implementation. Each model was fitted to responses generated by its own process (50 simulated agents per model). CBF recovery correlations were 0.95 for *µ*, 0.81 for *η*, and 0.95 for the softmax parameter *β*; RBPF recovery was 0.59 for its dynamics parameter and 0.72 for its softmax parameter. These results extend the recovery comparison to a different task and latent structure (Appendix F, Table 6).

## 5 Discussion

This paper introduced the CBF, a deterministic and differentiable alternative to particle-filter inference for sequential cognitive models with low-dimensional, bounded parameter spaces. Like prior particle-filter approaches to this problem, the CBF separates tractable conditional inference over latent states from intractable inference over noise parameters [11], but replaces the stochastic outer layer with a categorical distribution on a quantile grid. The procedure factors into exact discrete Bayesian inference under a parameterized prior and gradient-based optimization of the prior parameters, with the exact marginal likelihood serving as the optimization objective. On the volatility-stochasticity task published previously [11], fitted CBF dispersion parameters revealed a cross-over phenotyping pattern between volatility-blind and stochasticity-blind participants that is not recoverable from RBPF parameters fit to the same data even under a state-of-the-art protocol. The deterministic structure also surfaces a trial-by-trial computational ambiguity signal that predicts response times, a process-level analysis that stochastic methods can only approximate noisily.

### Implications for computational phenotyping

The cross-over result is consistent with the substantive claim that has motivated the dual-noise framework: that volatility-blindness and stochasticity-blindness are dimension-specific failure modes, not just two surface manifestations of the same underlying deficit [1]. The CBF makes this dimension-specific structure visible in the inferred priors. A subject behaviorally insensitive to volatility has a narrow prior over volatility values and a typical prior over stochasticity values; the converse holds for stochasticity-blindness. This kind of representation-level dissociation is exactly what computational phenotyping aims to deliver: not a single summary statistic of “how adaptive” a subject is, but a multidimensional account of which specific aspects of inference have gone awry. The RBPF, despite being fit to the same data, recovers the average behavioral pattern but does not reliably resolve the dimension-specific structure at the individual level. The methodological lesson is that the inferential apparatus matters as much as the conceptual model: even when two methods are designed to fit the same generative process, the one whose objective is differentiable and whose likelihood is deterministic delivers individual-level structure that the stochastic alternative cannot.

### Pathology lives in the extremes

The RBPF’s failure has a specific mechanism. The pathological groups are precisely those whose generative parameters lie near the boundary of the admissible range: a volatility-blind subject corresponds to a very low RBPF volatility update rate, a stochasticity-blind subject to a very low RBPF stochasticity update rate. Two compounding problems make these subjects hard to fit with a particle filter. First, the likelihood surface flattens near the boundary, since small variations in the parameter produce indistinguishable behavior and the data weakly constrain the parameter. Second, the particle filter likelihood estimator is noisier in the extreme regime, since the effective particle support contracts when most particles cluster around a value the parameter rarely visits. Together, these effects mean that the subjects whose parameters carry the most diagnostic information are estimated with the least precision. The CBF avoids both: the dispersion parameter *η* enters the likelihood through a smooth, differentiable parameterization that remains well-defined at the boundary, and the likelihood is deterministic, so the noise floor does not depend on the parameter regime.

### Broader methodological implication

The same logic generalizes beyond volatility-stochasticity inference. Particle filter models occupy an awkward position in cognitive science: conceptually appropriate for many problems involving inference over latent structure, but practically limited by stochastic likelihoods and non-differentiable objectives that force derivative-free fitting protocols and discourage individual-difference analyses where small effect sizes leave little room for estimator noise. The CBF resolves both limitations within the regime that covers most cognitive applications. The recipe is general: replace particles with a quantile grid, replace resampling with normalization, and parameterize the grid through differentiable quantile functions whose parameters can be optimized by gradient ascent. The cognitive-modeling literature contains many candidate applications, from conditioning [6] to category learning [5] to syntactic processing [9]. Our application of CBF to a social latent-cause task provides a demonstration of this broader applicability to distinct task domains and latent structures.

### Limitations

The CBF inherits its central limitation from the choice of representation. The categorical grid scales as *M* ^*D*^ in the parameter dimension, which is practical for small *D* but prohibitive beyond. This restricts the framework to problems where the unknown structure can be summarized in a small number of parameters, and excludes settings with many simultaneously varying latent factors or with structural parameters of high cardinality. The bounded-support assumption is similarly restrictive: the framework requires that admissible parameter values lie within a region selected a priori. The fixed grid also implicitly assumes static parameters within the inference horizon, with non-stationarity handled across blocks rather than online; a fully time-varying treatment would require either a moving grid or an additional layer of inference over change points.

The categorical-grid representation handles the regime relevant to most cognitive applications, and extensions to higher-dimensional problems, time-varying parameters, and unbounded supports are natural next steps. More broadly, the deterministic, differentiable structure makes hierarchical estimation, gradient-based fitting, and modern Bayesian workflows directly available for cognitive models that have so far been restricted to derivative-free protocols. The path from theoretical demonstration to practical inferential tool runs through this kind of methodological infrastructure.

## 6 Conclusion

Particle filters are a popular tool in cognitive modeling for inference over continuous latent parameters. A key limitation of particle filters, however, is that their likelihood is not differentiable with respect to the generative parameters of the cognitive model. Fitting therefore requires derivative-free methods, and the resulting parameter estimates are noisy enough to attenuate correlations between fitted parameters and other measures, which is the main use of such models in cognitive modeling and computational psychiatry. The CBF solves this problem: a categorical distribution over a differentiably parameterized quantile grid gives a marginal likelihood that is exact, deterministic, and differentiable in the generative parameters, so fitting becomes ordinary gradient-based evidence maximization. This is important because the gain is not only computational. In the volatility-stochasticity setting, the CBF recovered more of the behavioral signal than the particle filter, and its parameters separated maladaptation on one dimension from maladaptation on the other. The method is general for models with low-dimensional continuous latent parameters.

## Acknowledgments and Disclosure of Funding

This work was supported by an NIMH grant (R21MH134217) and a National Science Foundation CAREER Award (2543567) to Payam Piray. The authors declare no competing financial interests.

## A CBF fitting procedure

The CBF was fitted to behavioral data via Maximum A Posteriori (MAP) estimation. The procedure described here applies to both synthetic and human data; in both cases, fitting was performed independently for each individual dataset (a synthetic agent or a human subject). For each individual, four free parameters were estimated: the mean and dispersion of the Beta distribution for volatility (*µ*_*v*_, *η*_*v*_), and the mean and dispersion for stochasticity (*µ*_*s*_, *η*_*s*_), as defined in Section 3.3.

To ensure numerical stability and constrain the Beta distributions to valid ranges, the optimization was performed in an unconstrained parameter space (*x, y*) that was mapped to the model parameters via sigmoid transformations. For each dimension (volatility or stochasticity), the mean and dispersion were defined as

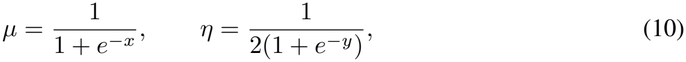

yielding *µ* ∈ (0, 1) and *η* ∈ (0, 0.5). The shape parameters of the Beta distribution are then *α* = *µ/η* and *β* = (1 − *µ*)*/η*, with *α* + *β* = 1*/η*. The constraint *η <* 0.5 guarantees *α* + *β >* 2, preventing U-shaped Beta configurations.

Because the CBF marginal likelihood is differentiable in the model parameters, exact gradients are available, and the negative log-posterior was minimized via standard gradient-based optimization. The model was fitted using the cbm toolbox [33], which uses Laplace approximation for model fitting and for approximating model evidence. Weakly informative Gaussian priors were placed on the unconstrained parameters: *x, y* ∼ *N* (0, 6.25), following the standard choice in cbm for parameters in the unit range. To mitigate sensitivity to initialization, the optimization was run from 10 random initializations per individual, and the best-converged solution was retained. This pulls the estimates toward the center of the sigmoid, providing mild regularization without strongly constraining the fit. We fixed the support of the volatility and stochasticity grids to [*v*_min_, *v*_max_] = [*s*_min_, *s*_max_] = [1, 70], and the number of atoms per dimension to *M* = 10. These quantities were treated as model hyperparameters and were not fit at the individual level.

## B Method validation on synthetic data

### B.1 Posterior dynamics

To illustrate the CBF’s posterior inference dynamics on synthetic data, we generated observation sequences under the same task structure as the human experiment: a 2 × 2 factorial design crossing two levels of volatility and stochasticity. For this illustrative simulation, the low and high levels were set symmetrically for the two noise dimensions, with true volatility set to low (*v* = 20) or high (*v* = 50), and true stochasticity set to low (*s* = 20) or high (*s* = 50).

The simulated agent was generated with *x*_vol_ = *x*_sto_ = 0 and *y*_vol_ = *y*_sto_ = 2. For this simulation, we fixed the volatility and stochasticity grid supports to [*v*_min_, *v*_max_] = [*s*_min_, *s*_max_] = [1, 70], with *M* = 20 atoms per dimension.

We generated 50 independent observation sequences for each condition and replayed the CBF on each sequence. Trajectories in Fig. 1 show the mean posterior estimate across runs, and shaded regions denote the standard error of the mean.

### B.2 Parameter recovery

The recovery analysis reported in the main text (Fig. 2) used *N*_sim_ = 100 synthetic agents with parameters *x* and *y* drawn uniformly from the logit-space range [−3, 3] (corresponding approximately to 0.05–0.95 for *µ* and 0.025–0.475 for *η*). The task structure matched the empirical experiment (4 blocks of 50 trials). Two analyses were performed using the fitting procedure described in Appendix A: a first using Maximum Likelihood Estimation with no regularization (which achieved Spearman *r* ≈ 1.0 across all four parameters), and a second using the empirical Gaussian priors *N* (0, 6.25) on the unconstrained parameters and minimizing the negative log-posterior with gradient-based optimization (which achieved Spearman *r >* 0.98 across all four parameters).

## C Application to human data

### C.1 Dataset

We applied the CBF to a previously published behavioral dataset [11], which describes the data collection procedure (including IRB approval and platform details) in full. The data come from *N* = 643 human participants.

Participants performed a predictive inference task (the “Bird Task”) designed to systematically dissociate volatility and stochasticity. On each trial, participants were asked to move a bucket to catch a bag of coins dropped by a hidden bird. The horizontal position of the bird (the latent state *x*_*t*_) evolved over trials according to a Gaussian random walk, while the position of the bag (the observation *o*_*t*_) was generated from the bird’s position with additive Gaussian noise.

The task used a stationary 2 × 2 factorial design, manipulating two sources of noise across four stable blocks. True volatility (*v*) was described to participants as the “speed” of the bird and controlled the variance of the latent random walk, with levels set to low (*v* = 4) or high (*v* = 49). True stochasticity (*s*) was described as the “wind” and controlled the variance of the observation noise, with levels set to low (*s* = 16) or high (*s* = 64). Each block consisted of 50 trials.

### C.2 Compute resources

All experiments were run on CPUs. Fitting to a single human subject’s data takes a few seconds. Total compute for all experiments is well under one CPU core-day.

## D Additional analyses

### D.1 Block-wise learning rate correlations

To assess whether the CBF captures individual differences in learning rates within each task condition, the fitted CBF model was replayed for each subject to obtain trial-by-trial learning rates, which were then averaged within each of the four blocks. In parallel, blockwise empirical learning rates were obtained from the model-agnostic regression used in the original study [11]. The main text reports the correlation between CBF-derived and empirical learning rates at baseline (averaged across blocks) and on the volatility and stochasticity main effects. Here, we additionally report the Spearman correlation across subjects within each of the four conditions (Table 1).

**Table 1:** CBF captures individual differences in block-wise learning rates. Spearman correlations between human and CBF-derived learning rates across *N* = 643 subjects.

| Condition | *ρ* | *p* |
|---|---|---|
| Low volatility, low stochasticity | 0.710 | < 0.001 |
| High volatility, low stochasticity | 0.809 | < 0.001 |
| Low volatility, high stochasticity | 0.785 | < 0.001 |
| High volatility, high stochasticity | 0.836 | < 0.001 |

### D.2 Response time

We tested whether trial-by-trial response times were related to the difficulty of assigning an observation to volatility versus stochasticity (Table 2). The CBF provides a natural measure of this difficulty through the spread of the likelihood across the volatility-stochasticity atom grid. When the likelihood is similar across many atoms, the observation does not clearly distinguish among competing noise explanations; when the likelihood is concentrated on a smaller subset of atoms, the source of noise is more identifiable.

For each subject, response times were regressed against a regressor quantifying per-trial ambiguity, defined as the standard deviation of likelihoods across all atoms. Several control regressors were also included to ensure that any effect was attributable to the regressor of interest: per-atom likelihood mean, squared prediction error, estimated volatility, and estimated stochasticity. Predictors were z-scored within subject; response times were kept in their original scale. The regression was fit separately for each subject, and group-level effects were tested using one-sample t tests on the subject-level coefficients. Likelihood standard deviation was significantly negatively related to response time, and the effect remained significant after controlling for the other predictors. Because response times were not used in fitting the CBF, this is a held-out behavioral signature of the model’s trial-level explaining-away computation.

**Table 2:** Response time regression results. *N* = 643 subjects.

| Predictor | *β* | *t*(642) | *p* |
|---|---|---|---|
| **Likelihood standard deviation** | **−8.93** | **−4.58** | **5.47 × 10<sup>−6</sup>** |
| Likelihood mean | 4.22 | 1.78 | 0.0759 |
| Squared prediction error | −0.48 | −0.31 | 0.7564 |
| Estimated volatility | −2.15 | −1.21 | 0.2259 |
| Estimated stochasticity | −3.95 | −2.15 | 0.0322 |

### D.2 Model comparison

We compared CBF and RBPF using random-effects Bayesian model selection across all subjects (Table 3). CBF was fitted separately by maximum likelihood in its native parameter space, without parameter priors, using *M* = 10 and volatility and stochasticity supports of [1, 70]. We used BIC-based approximations to log model evidence for both these CBF fits and the supplied RBPF fits from the original study [11]. The analysis took the subject-level model evidence values for each model and treated model identity as a random effect across subjects, estimating how frequently each model is expressed in the population [31, 32].

**Table 3:**
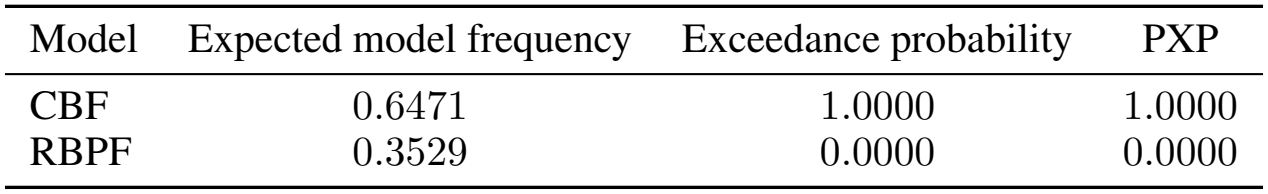
Bayesian model selection results. The Bayes omnibus risk was *<* 10^−6^. PXP = protected exceedance probability.

The expected model frequency was substantially higher for CBF than for RBPF (0.6471 versus 0.3529). The exceedance probability for CBF was 1.0000, indicating that CBF was overwhelmingly more likely than RBPF to have the larger population frequency. The protected exceedance probability (PXP) was also 1.0000, showing that this preference remains after accounting for the null possibility that the frequency differences arise by chance. The Bayes omnibus risk was below 10^−6^.

## E Robustness to grid resolution and support

We tested whether the cross-over depends on the number of atoms or the support of the volatility and stochasticity grids. We re-fitted the full sample of *N* = 643 subjects, varying the number of atoms per dimension over *M* ∈ *{*5, 10, 20} with the support upper bound fixed at 70, and varying the support upper bound over *v*_max_ = *s*_max_ ∈ {50, 70, 100} with *M* = 10. The lower bounds were fixed at *v*_min_ = *s*_min_ = 1. These are two separate sensitivity sweeps, sharing the standard *M* = 10, upper-bound-70 baseline.

At every setting, atom locations were constructed using the same quantile parameterization. The MAP fitting procedure and behavioral group definitions were otherwise unchanged. The volatility-blind (VB; *n* = 76) and stochasticity-blind (SB; *n* = 79) groups were held fixed across settings.

The cross-over persisted at every setting. Volatility-blind subjects had lower *η*_*v*_ than stochasticity-blind subjects, whereas stochasticity-blind subjects had lower *η*_*s*_ than volatility-blind subjects (Tables 4–5). Following the main analysis, group differences were tested using two-sided, independent-samples *t* tests on the unconstrained *y* coordinates, while the tables display transformed dispersion parameters. Both contrasts were significant at every setting (all *p <* 0.001, uncorrected; *df* = 153).

**Table 4:**
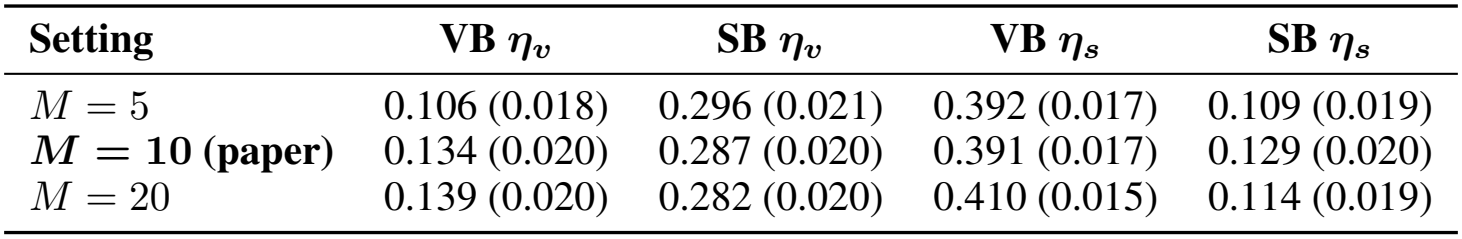
**Sensitivity to the number of atoms per dimension**, with *v*_max_ = *s*_max_ = 70. Entries are mean fitted dispersion (SEM). VB = volatility-blind; SB = stochasticity-blind.

| **Setting** | **VB *η*<sub>*v*</sub>** | **SB *η*<sub>*v*</sub>** | **VB *η*<sub>*s*</sub>** | **SB *η*<sub>*s*</sub>** |
|---|---|---|---|---|
| *M* = 5 | 0.106 (0.018) | 0.296 (0.021) | 0.392 (0.017) | 0.109 (0.019) |
| ***M* = 10 (paper)** | 0.134 (0.020) | 0.287 (0.020) | 0.391 (0.017) | 0.129 (0.020) |
| *M* = 20 | 0.139 (0.020) | 0.282 (0.020) | 0.410 (0.015) | 0.114 (0.019) |

**Table 5:**
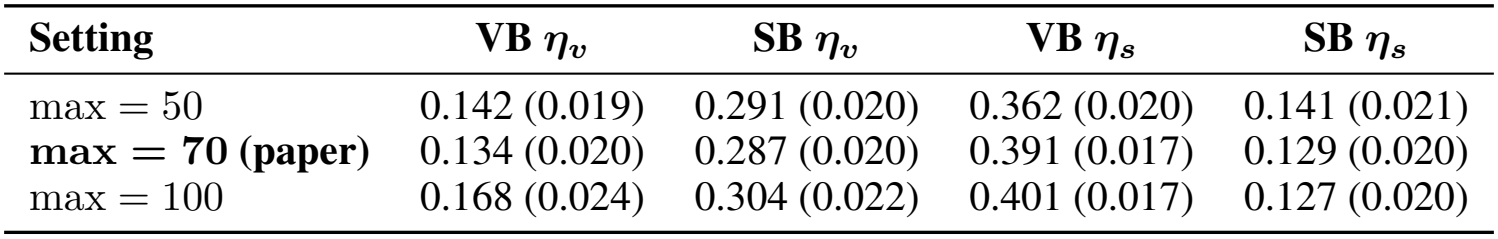
Sensitivity to the support upper bound, with *M* = 10 atoms per dimension. Entries are mean fitted dispersion (SEM). The lower bounds remain fixed at 1.

| **Setting** | **VB *η*<sub>*v*</sub>** | **SB *η*<sub>*v*</sub>** | **VB *η*<sub>*s*</sub>** | **SB *η*<sub>*s*</sub>** |
|---|---|---|---|---|
| max = 50 | 0.142 (0.019) | 0.291 (0.020) | 0.362 (0.020) | 0.141 (0.021) |
| **max = 70 (paper)** | 0.134 (0.020) | 0.287 (0.020) | 0.391 (0.017) | 0.129 (0.020) |
| max = 100 | 0.168 (0.024) | 0.304 (0.022) | 0.401 (0.017) | 0.127 (0.020) |

**Table 6:** Parameter self-recovery in the social latent-cause inference task. Spearman correlations between true and recovered parameters for 50 simulated subjects per model. Each model was fitted to data generated by its own process.

| Model | Dynamics | β (softmax) |
|---|---|---|
| CBF | μ = 0.95, η = 0.81 | 0.95 |
| RBPF | 0.59 | 0.72 |

**Table 7:** Volatility-blind (*n* = 76) vs. rest (*n* = 567)

| Measure | Group | Mean (SEM) | Median [25%, 75%] | Min–Max |
|---|---|---|---|---|
| *μ*<sub>*v*</sub> | Vol-blind | 0.393 (0.040) | 0.253 [0.092, 0.663] | 0.000–0.999 |
| *μ*<sub>*v*</sub> | Rest | 0.319 (0.013) | 0.174 [0.059, 0.509] | 0.000–0.995 |
| *μ*<sub>*s*</sub> | Vol-blind | 0.373 (0.045) | 0.122 [0.030, 0.794] | 0.000–0.999 |
| *μ*<sub>*s*</sub> | Rest | 0.298 (0.015) | 0.108 [0.034, 0.562] | 0.000–1.000 |

**Table 8:**
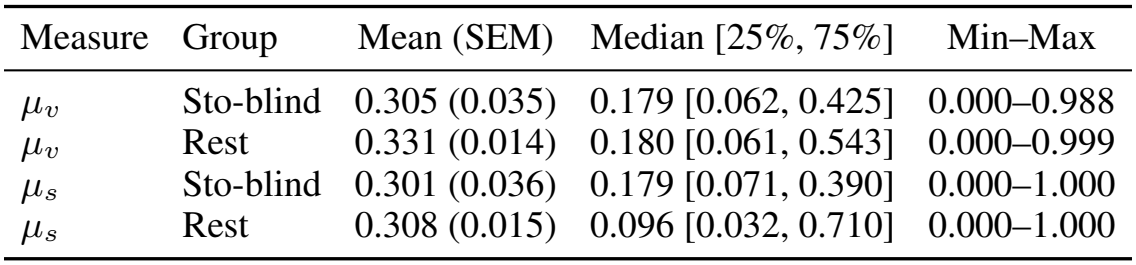
Stochasticity-blind (*n* = 79) vs. rest (*n* = 564)

| Measure | Group | Mean (SEM) | Median [25%, 75%] | Min–Max |
|---|---|---|---|---|
| *μ<sub>v</sub>* | Sto-blind | 0.305 (0.035) | 0.179 [0.062, 0.425] | 0.000–0.988 |
| *μ<sub>v</sub>* | Rest | 0.331 (0.014) | 0.180 [0.061, 0.543] | 0.000–0.999 |
| *μ<sub>s</sub>* | Sto-blind | 0.301 (0.036) | 0.179 [0.071, 0.390] | 0.000–1.000 |
| *μ<sub>s</sub>* | Rest | 0.308 (0.015) | 0.096 [0.032, 0.710] | 0.000–1.000 |

**Table 9:**
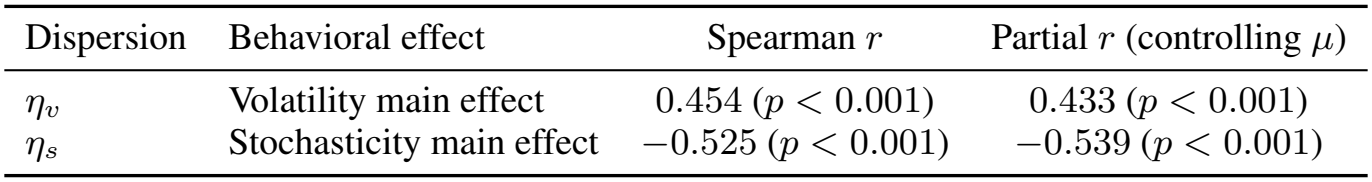
Associations between *η*_*v*_ or *η*_*s*_ and the corresponding behavioral main effect across *N* = 643 participants without behavioral grouping. Partial Spearman correlations control for *µ*_*v*_ when testing *η*_*v*_, and for *µ*_*s*_ when testing *η*_*s*_.

| Dispersion | Behavioral effect | Spearman *r* | Partial *r* (controlling *μ*) |
|---|---|---|---|
| *η<sub>v</sub>* | Volatility main effect | 0.454 (*p* < 0.001) | 0.433 (*p* < 0.001) |
| *η<sub>s</sub>* | Stochasticity main effect | −0.525 (*p* < 0.001) | −0.539 (*p* < 0.001) |

**Table 10:** Associations between psychiatric dimensions and model parameters.

| Source | Measure | *r* | *p* |
|---|---|---|---|
| CBF | ***η<sub>s</sub>*** | **−0.157** | **0.0016** |
| CBF | *η<sub>v</sub>* | −0.028 | 0.57 |
| CBF | *μ<sub>v</sub>* / *μ<sub>s</sub>* | −0.032 / −0.002 | 0.52 / 0.97 |
| RBPF | Volatility / stochasticity update rate | −0.034 / −0.003 | 0.50 / 0.96 |
| Model-neutral | Volatility / stochasticity effect | +0.051 / +0.017 | 0.31 / 0.74 |

Thus, the qualitative phenotyping result was not dependent on the tested atom resolution or upper bound.

Increasing *M* from 10 to 20 changed group-mean dispersion estimates by approximately 2–12% relative to their values at *M* = 10 while preserving the cross-over. Even *M* = 5 (25 joint atoms) retained both group contrasts, so the result is not a fine-resolution artifact.

## F Generalization to a social latent-cause inference task

The reliability gap between CBF and RBPF replicates on a second task with a different RBPF implementation. We applied the CBF to a social latent-cause inference task [14], in which participants attribute jointly produced outcomes to one of three agents presented in pairs. The task differs from the Bird Task on every relevant axis: the latent cause is a discrete responsibility assignment (pairwise responsibilities *z*_12_, *z*_13_, *z*_23_ over shared abilities) rather than a continuous volatility/stochasticity split, with three interacting sources rather than a 2 × 2 factorial, in a social rather than perceptual domain; and the baseline is a different RBPF instance (Rao-Blackwellized over discrete responsibility). Self-recovery (each model fit to data generated by its own process; *n* = 50 simulated subjects), with Spearman correlations between true and recovered parameters:

(CBF notation as in the main text: grid mean *µ* and dispersion *η* over the dynamics parameter. The RBPF’s dynamics parameter is a diffusion probability, i.e., the probability that a binary responsibility flips, a different quantity, so we leave it unsymbolized to avoid collision with the CBF *µ*.)

The CBF attains near-ceiling self-recovery of the parameters it generated; RBPF self-recovery is substantially noisier (*r* = 0.59 for its dynamics parameter), closely echoing the reliability gap on the Bird Task (*r* = 0.69*/*0.67; Sec. 4.1). Notably, this task has a separate, dynamics-independent softmax *β* parameter shared by both models; RBPF recovery of this parameter is also much noisier than the CBF’s (RBPF: *r* = 0.72 vs. CBF: *r* = 0.95). The identifiability advantage of our approach is therefore not specific to one task, one model structure, or one RBPF implementation.

## G Controls for boundary effects and joint parameter fitting

### G.1 Distributions of *µ*_*v*_ and *µ*_*s*_ across behavioral groups

Because *µ* and *η* jointly determine the Beta distribution, we examined whether the group differences in *η*_*v*_ and *η*_*s*_ were accompanied by differences in *µ*_*v*_ and *µ*_*s*_. In particular, we assessed whether *µ* estimates were concentrated near the parameter boundaries and whether the dispersion–behavior associations remained after controlling for the corresponding *µ* parameter.

We first compared the fitted *µ*_*v*_ and *µ*_*s*_ distributions in each blind group with those of the remaining participants.

The fitted *µ* distributions were broad and overlapped those of the remaining participants. These descriptive summaries do not indicate that the blind groups were confined to a narrow region near the parameter boundaries.

### G.2 Associations between *η* and behavior remain significant after controlling for *µ*

We next tested whether the associations between *η* and behavior extended across the full sample rather than depending on the behavioral group thresholds. Across all *N* = 643 participants, we computed Spearman correlations between *η*_*v*_ and the empirical volatility main effect, and between *η*_*s*_ and the empirical stochasticity main effect, without thresholding or group labels. To assess whether these associations were attributable to the corresponding *µ* parameter, we also computed partial Spearman correlations controlling for *µ*_*v*_ or *µ*_*s*_ respectively.

Each dispersion parameter tracked its corresponding behavioral dimension, and both associations remained significant after controlling for the corresponding *µ* parameter. The off-dimension association is null (*η*_*v*_ with the stochasticity effect: *r* = − 0.005, *p* = 0.905). Because these analyses use the full sample without behavioral group labels, the associations do not depend on the classify-then-compare procedure used to visualize the cross-over in Fig. 3. The group comparison illustrates this continuous relationship. As a further comparison, the RBPF fitted to the identical groups does not reproduce the cross-over (Fig. 3, right).

Together, these controls support the interpretation that the cross-over reflects dimension-specific associations between *η* and behavior, rather than merely boundary-concentrated *µ* estimates or associations explained by the corresponding fitted *µ* parameter.

## H Associations with psychiatric questionnaire measures

We examined whether individual differences in fitted CBF parameters were associated with psychiatric symptom measures collected after the learning task. These measures provide an external assessment of the potential relevance of the fitted parameters beyond the behavioral quantities used for model fitting and group classification.

Our *N* = 643 sample combines the two datasets reported in [11] (*n* = 223 and *n* = 420); the psychiatric battery analyzed here was administered in the second (*n* = 401 after attention-check exclusions). These questionnaire data were collected alongside the behavioral task but not analyzed in [11], and they enter neither the fitting nor the classification. The battery comprises the PSWQ-8 (worry) and the 7 Up 7 Down (yielding separate hypomania and depression scores), giving three symptom scores per participant; the GAD-7 was also collected but is not analyzed here. The framework of Piray and Daw [1] predicts that a low stochasticity dispersion (*η*_*s*_) is the signature associated with the intolerance of uncertainty and chronic worry characteristic of anxiety.

We proceeded in two steps. First, each fitted parameter was regressed on a design matrix containing all three scores. *η*_*s*_ was the only parameter showing any association: negatively with worry (*b* = − 0.79, *p* = 0.0018) and positively with depression (*b* = +0.65, *p* = 0.011), with hypomania null (*p* = 0.40); no other CBF parameter, *η*_*v*_ included, showed any association (all *p >* 0.10). Second, because worry and depression are collinear in this sample (*r* = 0.67), their separate coefficients are mutually suppressed and not individually interpretable; the opposite signs are consistent with *η*_*s*_ tracking their difference, so we formalized this as the standardized contrast *z*(worry) − *z*(depression) and tested it against every parameter:

*η*_*s*_ is the only parameter with an association, surviving Bonferroni correction across the four CBF parameters (*p* = 0.006), and its direction, lower *η*_*s*_ accompanying a symptom profile weighted toward worry rather than depressed mood, is the one predicted. The worry-minus-depression contrast was defined after inspecting the collinearity between the two scores.

## I Code and data availability

The implementation, fitting procedures, and analysis scripts are available at https://github.com/piraylab/CBF. The repository includes instructions for fitting models from scratch and reproducing analyses using saved fits.

### NeurIPS Paper Checklist

1. **Claims** Question: Do the main claims made in the abstract and introduction accurately reflect the paper’s contributions and scope? Answer: [Yes] Justification: The abstract’s claims are supported by theoretical and empirical evidence (Sections 3 and 4).
2. **Limitations** Question: Does the paper discuss the limitations of the work performed by the authors? Answer: [Yes] Justification: A paragraph of Section 5 discusses limitations.
3. **Theory assumptions and proofs** Question: For each theoretical result, does the paper provide the full set of assumptions and a complete (and correct) proof? Answer: [Yes] Justification: We provide complete proofs or derivation for our theoretical results in Section 3.
4. **Experimental result reproducibility** Question: Does the paper fully disclose all the information needed to reproduce the main experimental results of the paper to the extent that it affects the main claims and/or conclusions of the paper (regardless of whether the code and data are provided or not)? Answer: [Yes] Justification: We provide all details of our algorithms in Section 3 and Appendix A.
5. **Open access to data and code** Question: Does the paper provide open access to the data and code, with sufficient instructions to faithfully reproduce the main experimental results, as described in supplemental material? Answer: [Yes] Justification: Code, the public behavioral datasets, and instructions for model fitting and analysis are available at https://github.com/piraylab/CBF. The repository documents how to fit models from scratch and rerun analyses using saved fits, and identifies any reported analyses whose supporting code and data are not included.
6. **Experimental setting/details** Question: Does the paper specify all the training and test details (e.g., data splits, hyperparameters, how they were chosen, type of optimizer) necessary to understand the results? Answer: [Yes] Justification: The main text provides experimental settings sufficient for understanding, with additional details, including hyperparameters and fitting procedures, in Appendix A.
7. **Experiment statistical significance** Question: Does the paper report error bars suitably and correctly defined or other appropriate information about the statistical significance of the experiments? Answer: [Yes] Justification: We report mean and standard error of the mean wherever applicable.
8. **Experiments compute resources** Question: For each experiment, does the paper provide sufficient information on the computer resources (type of compute workers, memory, time of execution) needed to reproduce the experiments? Answer: [Yes] Justification: We discuss compute resources in Appendix C.
9. **Code of ethics** Question: Does the research conducted in the paper conform, in every respect, with the NeurIPS Code of Ethics https://neurips.cc/public/EthicsGuidelines? Answer: [Yes] Justification: This paper conforms with the NeurIPS Code of Ethics.
10. **Broader impacts** Question: Does the paper discuss both potential positive societal impacts and negative societal impacts of the work performed? Answer: [Yes] Justification: The paper’s potential positive impact is discussed implicitly throughout: the proposed method enables more reliable individual-difference and clinical-trait analyses in cognitive modeling and computational psychiatry. The work does not introduce new data, models of human behavior, or systems with foreseeable misuse potential beyond those already standard in the field, and we do not foresee specific negative societal impacts arising from the methodological contribution.
11. **Safeguards** Question: Does the paper describe safeguards that have been put in place for responsible release of data or models that have a high risk for misuse (e.g., pre-trained language models, image generators, or scraped datasets)? Answer: [N/A] Justification: This paper poses no such risks.
12. **Licenses for existing assets** Question: Are the creators or original owners of assets (e.g., code, data, models), used in the paper, properly credited and are the license and terms of use explicitly mentioned and properly respected? Answer: [Yes] Justification: All original owners of the data used in the paper are properly credited.
13. **New assets** Question: Are new assets introduced in the paper well documented and is the documentation provided alongside the assets? Answer: [N/A] Justification: The paper does not introduce new datasets, pre-trained models, or software libraries as released assets. The new method is fully specified in Section 3.
14. **Crowdsourcing and research with human subjects** Question: For crowdsourcing experiments and research with human subjects, does the paper include the full text of instructions given to participants and screenshots, if applicable, as well as details about compensation (if any)? Answer: [N/A] Justification: This paper reanalyzes a previously published behavioral dataset [11]; no new human subjects data was collected. Task instructions, compensation, and ethics approval are described in the cited original paper.
15. **Institutional review board (IRB) approvals or equivalent for research with human subjects** Question: Does the paper describe potential risks incurred by study participants, whether such risks were disclosed to the subjects, and whether Institutional Review Board (IRB) approvals (or an equivalent approval/review based on the requirements of your country or institution) were obtained? Answer: [N/A] Justification: This paper reanalyzes a previously published behavioral dataset [11]; no new human subjects data was collected. Ethics approval is described in the cited original paper.
16. **Declaration of LLM usage** Question: Does the paper describe the usage of LLMs if it is an important, original, or non-standard component of the core methods in this research? Note that if the LLM is used only for writing, editing, or formatting purposes and does *not* impact the core methodology, scientific rigor, or originality of the research, declaration is not required. Answer: [N/A] Justification: LLMs have only been used for drafting, editing, formatting, and coding assistance.

## References

[1] Payam Piray and Nathaniel D. Daw. A model for learning based on the joint estimation of stochasticity and volatility. Nature Communications, 12(1):6587, November 2021. ISSN 2041-1723. doi:10.1038/s41467-021-26731-9. URL https://www.nature.com/articles/s41467-021-26731-9.

[2] Timothy E J Behrens, Mark W Woolrich, Mark E Walton, and Matthew F S Rushworth. Learning the value of information in an uncertain world. Nature Neuroscience, 10(9):1214–1221, September 2007. ISSN 1097-6256, 1546-1726. doi:10.1038/nn1954. URL https://www.nature.com/articles/nn1954.

[3] Christoph Mathys. A Bayesian foundation for individual learning under uncertainty. Frontiers in Human Neuroscience, 5, 2011. ISSN 16625161. doi:10.3389/fnhum.2011.00039. URL http://journal.frontiersin.org/article/10.3389/fnhum.2011.00039/abstract.

[4] Payam Piray and Nathaniel D. Daw. A simple model for learning in volatile environments. PLOS Computational Biology, 16(7):e1007963, July 2020. ISSN 1553-7358. doi:10.1371/journal.pcbi.1007963. URL https://dx.plos.org/10.1371/journal.pcbi.1007963.

[5] Adam N. Sanborn, Thomas L. Griffiths, and Daniel J. Navarro. Rational approximations to rational models: Alternative algorithms for category learning. Psychological Review, 117(4):1144–1167, 2010. ISSN 1939-1471, 0033-295X. doi:10.1037/a0020511. URL https://doi.apa.org/doi/10.1037/a0020511.

[6] Nathaniel Daw and Aaron Courville. The pigeon as particle filter. Advances in neural information processing systems, 20:369–376, 2008.

[7] Samuel J Gershman, David M Blei, and Yael Niv. Context, learning, and extinction. Psychological review, 117(1):197, 2010.

[8] Scott D Brown and Mark Steyvers. Detecting and predicting changes. Cognitive psychology, 58(1):49–67, 2009.

[9] Roger Levy. Expectation-based syntactic comprehension. Cognition, 106(3):1126–1177, 2008.

[10] C Maher, I Saez, and A Radulescu. Decoding covert human attention in multidimensional environments. bioRxiv, pages 2026–03, 2026.

[11] Payam Piray and Nathaniel D. Daw. Computational processes of simultaneous learning of stochasticity and volatility in humans. Nature Communications, 15(1):9073, October 2024. ISSN 2041-1723. doi:10.1038/s41467-024-53459-z. URL https://www.nature.com/articles/s41467-024-53459-z.

[12] Xiaotong Fang and Payam Piray. Inferring the causes of noise from binary outcomes: A normative theory of learning under uncertainty. bioRxiv, pages 2026–03, 2026.

[13] Marc G Bellemare, Will Dabney, and Rémi Munos. A distributional perspective on reinforcement learning. In International conference on machine learning, pages 449–458. Pmlr, 2017.

[14] Ali Mahmoodi, Shuyi Luo, Caroline Harbison, Payam Piray, and Matthew F. S. Rushworth. Human hippocampus and dorsomedial prefrontal cortex infer and update latent causes during social interaction. Neuron, 112(22):3796–3809.e9,2024. doi: 10.1016/j.neuron.2024.09.001.

[15] Angela J. Yu and Peter Dayan. Uncertainty, Neuromodulation, and Attention. Neuron, 46(4):681–692, May 2005. ISSN 08966273. doi:10.1016/j.neuron.2005.04.026. URL https://linkinghub.elsevier.com/retrieve/pii/S0896627305003624.

[16] Alireza Soltani and Alicia Izquierdo. Adaptive learning under expected and unexpected uncertainty. Nature Reviews Neuroscience, 20(10):635–644, 2019.

[17] Matthew R. Nassar, Robert C. Wilson, Benjamin Heasly, and Joshua I. Gold. An Approximately Bayesian Delta-Rule Model Explains the Dynamics of Belief Updating in a Changing Environment. The Journal of Neuroscience, 30(37):12366–12378, September 2010. ISSN 0270-6474, 1529-2401. doi:10.1523/JNEUROSCI.0822-10.2010. URL https://www.jneurosci.org/lookup/doi/10.1523/JNEUROSCI.0822-10.2010.

[18] Albert R Powers, Christoph Mathys, and Philip Robert Corlett. Pavlovian conditioning–induced hallucinations result from overweighting of perceptual priors. Science, 357(6351):596–600, 2017.

[19] Rebecca P Lawson, Christoph Mathys, and Geraint Rees. Adults with autism overestimate the volatility of the sensory environment. Nature neuroscience, 20(9):1293–1299, 2017.

[20] Kevin Murphy and Stuart Russell. Rao-blackwellised particle filtering for dynamic bayesian networks. In Sequential Monte Carlo methods in practice, pages 499–515. Springer, 2001.

[21] Christian Naesseth, Scott Linderman, Rajesh Ranganath, and David Blei. Variational sequential monte carlo. In International conference on artificial intelligence and statistics, pages 968–977. PMLR, 2018.

[22] Tuan Anh Le, Maximilian Igl, Tom Rainforth, Tom Jin, and Frank Wood. Auto-encoding sequential monte carlo. arXiv preprint arXiv:1705.10306, 2017.

[23] Chris J Maddison, John Lawson, George Tucker, Nicolas Heess, Mohammad Norouzi, Andriy Mnih, Arnaud Doucet, and Yee Teh. Filtering variational objectives. Advances in neural information processing systems, 30, 2017.

[24] Rico Jonschkowski, Divyam Rastogi, and Oliver Brock. Differentiable particle filters: End-to-end learning with algorithmic priors. arXiv preprint arXiv:1805.11122, 2018.

[25] Adrien Corenflos, James Thornton, George Deligiannidis, and Arnaud Doucet. Differentiable particle filtering via entropy-regularized optimal transport. In International Conference on Machine Learning, pages 2100–2111. PMLR, 2021.

[26] Ardavan Saeedi, Tejas D Kulkarni, Vikash K Mansinghka, and Samuel J Gershman. Variational particle approximations. Journal of Machine Learning Research, 18(69):1–29, 2017.

[27] Will Dabney, Mark Rowland, Marc G. Bellemare, and Rémi Munos. Distributional Reinforcement Learning with Quantile Regression, October 2017. URL http://arxiv.org/abs/1710.10044. arXiv:1710.10044 [cs].

[28] Will Dabney, Zeb Kurth-Nelson, Naoshige Uchida, Clara Kwon Starkweather, Demis Hassabis, Rémi Munos, and Matthew Botvinick. A distributional code for value in dopamine-based reinforcement learning. Nature, 577(7792):671–675, January 2020. ISSN 0028-0836, 1476-4687. doi:10.1038/s41586-019-1924-6. URL https://www.nature.com/articles/s41586-019-1924-6.

[29] Genshiro Kitagawa. Non-gaussian state—space modeling of nonstationary time series. Journal of the American statistical association, 82(400):1032–1041, 1987.

[30] Richard S Bucy and Kenneth D Senne. Digital synthesis of non-linear filters. Automatica, 7(3):287–298, 1971.

[31] Klaas Enno Stephan, Will D Penny, Jean Daunizeau, Rosalyn J Moran, and Karl J Friston. Bayesian model selection for group studies. Neuroimage, 46(4):1004–1017, 2009.

[32] Lionel Rigoux, Klaas Enno Stephan, Karl J Friston, and Jean Daunizeau. Bayesian model selection for group studies—revisited. Neuroimage, 84:971–985, 2014.

[33] Payam Piray, Amir Dezfouli, Tom Heskes, Michael J Frank, and Nathaniel D Daw. Hierarchical bayesian inference for concurrent model fitting and comparison for group studies. PLoS computational biology, 15 (6):e1007043, 2019.

